# Oropouche virus results in severe congenital disease in embryonic mice

**DOI:** 10.1101/2025.09.23.678046

**Authors:** Maïlis Darmuzey, Robert M. Verdijk, Niels Cremers, Martin Ferrié, Stijn Hendrickx, Alexandre Hego, Suzanne J. F. Kaptein, Johan Neyts

**Affiliations:** KU Leuven Department of Microbiology, Immunology and Transplantation, Virology, Antiviral Drug & Vaccine Research Group, Rega Institute for Medical Research, Leuven, Belgium; Department of Pathology, Section Ophthalmic Pathology, Erasmus MC University Medical Center, Rotterdam, The Netherlands; Cell Imaging Platform, GIGA Institute, University of Liège, Belgium

## Abstract

The recent outbreak of Oropouche virus (OROV) in Latin-America with unprecedented reports of vertical transmission resulting in congenital malformations and fetal death, redefines the threat posed by this arbovirus. We here explore the teratogenic potential of an epidemic strain of the virus in a mouse model, whereby a low inoculum of OROV is injected into the placenta. The virus replicates efficiently in multiple embryonic organs, with high titers in the brain, and causes intrauterine growth restriction. Infected embryos display multi-organ pathology, including pneumonitis, myocarditis, steatohepatitis and severe neuropathology. Embryonic brains exhibit microcephaly, ventriculomegaly and extensive neural loss, mirroring findings in human fetal cases. A dramatic alteration of the neural cell population correlates with massive cell death, demonstrating the extreme cytotoxicity of OROV for the developing brain. The infection also triggers neuroinflammation characterized by cytokine upregulation, microglial activation and neutrophil infiltration. Altogether, our findings establish a link between congenital OROV infection and neurodevelopmental disease, highlighting its teratogenic potential.

## Main

Oropouche virus (OROV) is an arthropod-borne virus from the *Orthobunyavirus* genus, whose principal vector is the biting midge *Culicoides paraensis*^1^. OROV is endemic in Central and South America^1^ and has been responsible for multiple sporadic epidemic events since its first isolation and identification in 1955 in Trinidad and Tobago^2^. Historically, OROV infections typically led to asymptomatic cases or nonspecific symptoms referred to as Oropouche fever, which is characterized by headache, arthralgia, myalgia, nausea, and vomiting, amongst others^3^. The small-scale Oropouche epidemics and the moderate and transient health issues caused by OROV infections led to the perception that OROV did not pose a major threat to public health. However, the 2023-2024 outbreak in Latin America marked an important turning point in light of its unprecedented magnitude, the exponential geographic spread to new countries and its impact on human health. For the first time, OROV infections were linked to fatal outcomes in infected individuals in the Americas^4^. Moreover, the Pan American Health Organization (PAHO) issued an epidemiological alert regarding possible cases of vertical transmission of OROV^5^. In 2024, five cases of OROV vertical transmission were confirmed in Brazil, with four cases of fetal death and the fifth case presenting with a congenital anomaly. Several cases, of which 22 of *in utero* fetal death and 4 cases of congenital defects, were additionally under investigation^4^. A recent study of 68 historical cases of congenital malformations identified six OROV-positive cases, including one case with severe brain defects with necrotic and apoptotic neurons, microglia and astrocytes, vacuolization and tissue atrophy. Viral RNA was detected in the brain, lung, kidney and cerebral spinal fluid, and viral antigens in brain, liver, kidney, heart and lungs tissues^6^. Altogether, the 2023-2024 OROV outbreak highlighted the threat posed by this virus to human health and to the developing fetus in particular.

Although it is recognized that vertical transmission of OROV and the associated miscarriages may result from an OROV infection, the underlying pathophysiology remains unclear. Moreover, although suspected, the link between vertical transmission and congenital disease resulting from the teratogenic potential of OROV has not been unequivocally established. Hence, there is an urgent need for a better understanding of the teratogenic potential of OROV in order to assess the potential risk of infection with this virus to the developing fetus. Here we examined the pathophysiology of an OROV infection after vertical transmission by via intraplacental (IPL) infection in mice. We demonstrated earlier the physiological relevance of this model for the study of Zika virus-induced congenital disease^7^. Given the alarming rise in OROV-associated congenital anomalies and fetal deaths, a deeper understanding of its pathogenic potential is imperative. Our study aims to shed light on the pathogenicity of OROV, which may compromise fetal development.

## Results

### OROV replicates efficiently in embryonic organs and causes intra-uterine growth retardation

To determine whether OROV can affect embryonic development, we performed IPL infections in the embryonic labyrinth with 10-20 plaque-forming units (PFU) of the epidemic OROV_IRCCS-_ ^8^ strain at embryonic day 12.5 (E12.5) (Fig. 1a). Five days post-infection (d5 pi), i.e. at E17.5, the effect on embryonic development was assessed by comparing the crown-rump length and weight of the infected embryos with those of their uninfected counterparts. We found that OROV-infected embryos manifested significant intra-uterine growth retardation, as evidenced by a smaller crown-rump length (Fig. 1b) and a lower body weight (Fig. 1c) compared to mock-infected embryos at the same developmental stage. When comparing the replication kinetics of OROV over the course of infection until E17.5 (d5 pi) in the different embryonic tissues (brain, lungs, heart and liver), a similar replication pattern was noted, i.e. the viral RNA load increased from E13.5 (d1 pi) to E16.5 (d4 pi; peak of replication), followed by a decrease on E17.5 (d5 pi) (Fig. 1d). However, while viral RNA levels dropped markedly in the lungs, heart and liver on E17.5, only a minor decrease was observed in the brains of embryonic mice. Moreover, notable differences were found between viral RNA levels in the embryonic brains and those in the other tissues examined on both E16.5 and E17.5. Whereas viral RNA levels were comparable in lungs, heart and liver on E16.5 (2.82E+08 averaged) and E17.5 (3.24E+06 averaged) levels in the brain were on average 4.65E+09 and 9.77E+08 on E16.5 and E17.5, respectively.

**Fig. 1:**
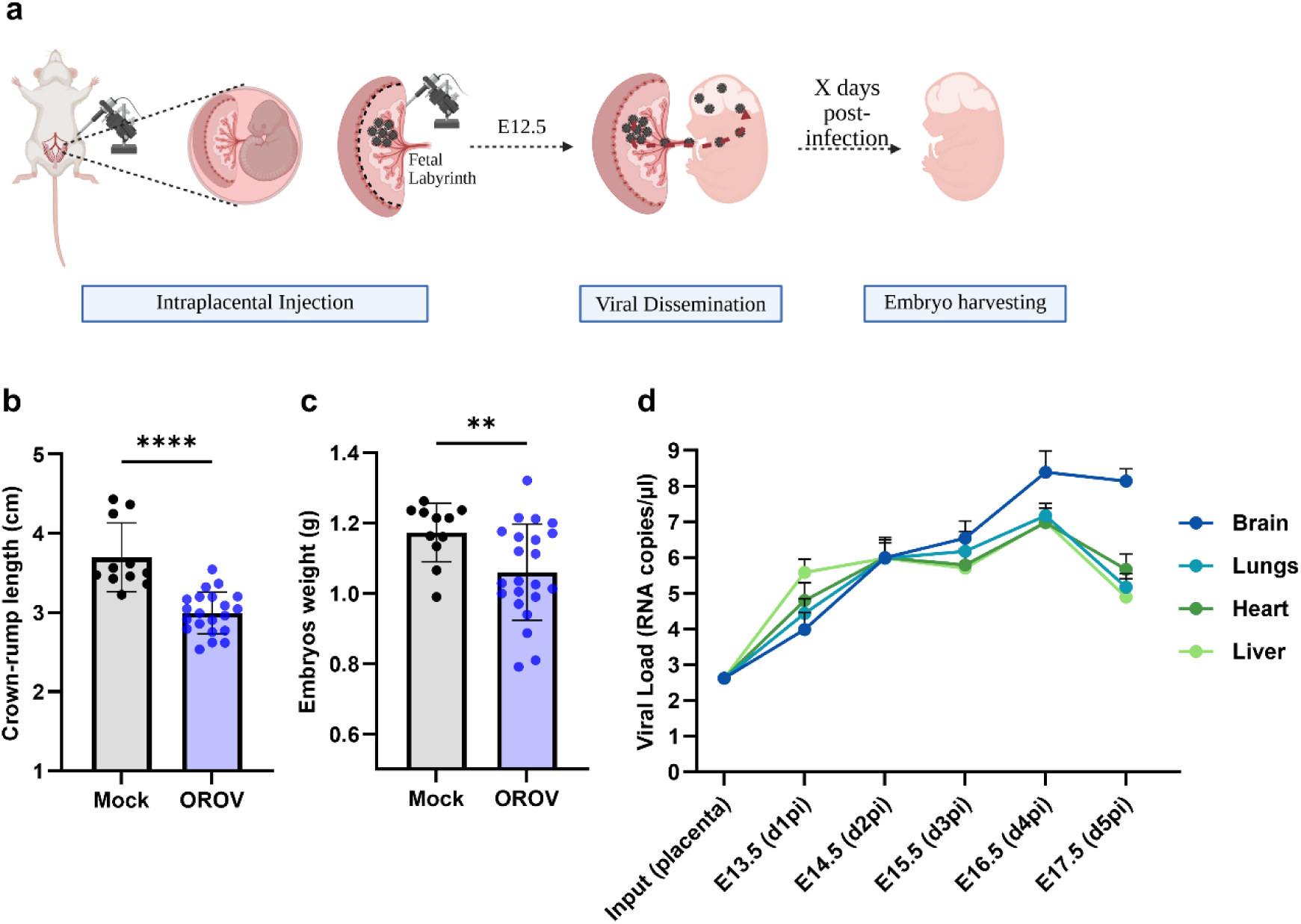
Congenital OROV infection causes IUGR and replicates in embryonic organs. **a** Schematic representation of the intraplacental infection with OROV at E12.5. **b** Measure of the crown-rump length of mock-infected and OROV-infected embryos at E17.5. **c** Measure of the weight of mock-infected and OROV-infected embryos at E17.5. **d** Replication kinetics of OROV in the embryonic brains, lungs, hearts and liver from E13.5 to E17.5. N numbers are shown in Supplementary Table 3. **b**,**c**, data are presented as mean ± standard deviation. Statistical significance of differences was determined by the unpaired t test with Welch’s correction (**b,c**). **d,** data are presented as mean ± S.E.M. Only statistically significant differences are shown (****p < 0.0001; **p < 0.01).

### OROV infection in embryonic mice induces severe multi-organ pathology

To further investigate the pathology caused by OROV in different organs (brain, spinal cord, lung, heart and liver), embryonic mice were intraplacentally infected at E12.5 and harvested at E17.5 (d5 pi). Entire embryos were subsequently processed by cryo-sectioning, stained with hematoxylin and eosin, and examined for abnormalities. The affected embryos typically exhibited delayed maturation consistent with the intrauterine growth retardation, and also presented some degree of hydrops (Supplementary Fig. 1). In line with the highest viral RNA load in the brain, the most severe damage was also seen in the brain, which presented severe necrotizing meningoencephalitis and hemorrhages (Fig. 2b, d). Besides the brain, the spinal cord also displayed a severe phenotype as demonstrated by the presence of hemorrhages and necrosis with mixed inflammatory infiltrations (Fig. 2f; Supplementary Fig. 2b and Supplementary Fig. 3a). Other organs of the embryonic mice, including the lungs, heart and liver, were also affected. More specifically, the lungs contained mixed interstitial and focal intra-alveolar infiltrates compatible with a most prominent pneumonitis and focal pneumonia (Fig. 2h), while the heart muscle was infiltrated with a mixed inflammatory cell population indicating a moderate myocarditis (Fig. 2j). The liver showed severe hemorrhagic steatohepatitis, as proven by a positive oil red O staining indicating the presence of lipids (Fig. 2l, n). The immune infiltrate in the internal organs consisted mostly of mononuclear cells with extensive admixture of neutrophils (i.e. Ly6G-positive cells) (Supplementary Fig. 2b, d), especially in the necrotic areas of the CNS. Taken together, the severe lesions in the brain and spinal cord indicate neurotropism. In addition, the hydrops as well as the overall delay in organ development and the multi-organ pathology point to a strong teratogenic potential of this virus.

**Fig. 2:**
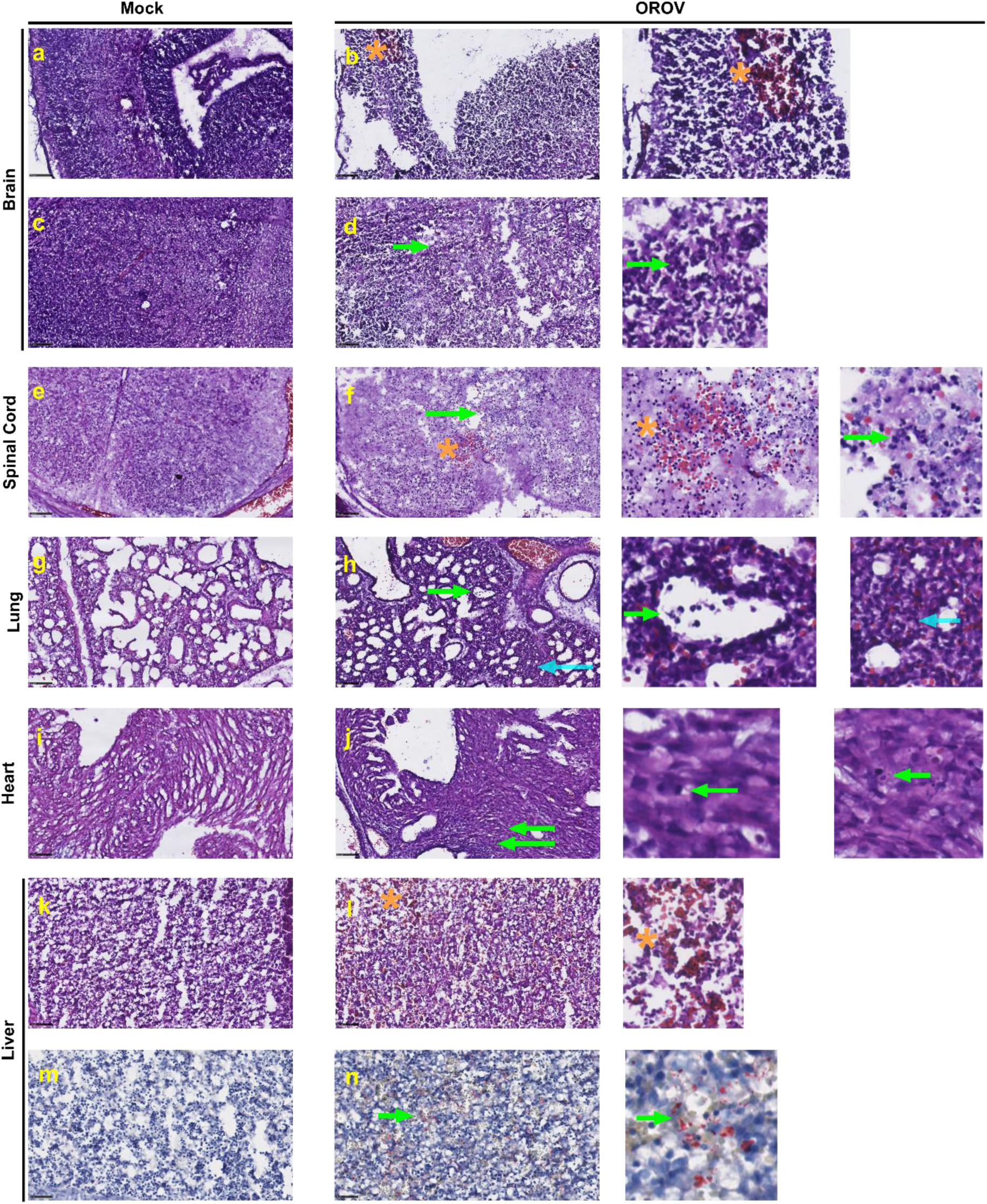
Congenital OROV infection causes severe pathological alteration in embryonic organs. **a-n** Representative hematoxylin and eosin-stained images at E17.5. **a-d** Representative image of mock-infected (**a,c**) and OROV-infected (**b,d**) brains, orange star and green arrow indicated hemorrhages and necrosis respectively. **e-f** Representative image of mock-infected (**e**) OROV-infected (**f**) spinal cord, orange star and green arrow indicated respectively hemorrhages and inflammatory infiltration. **g-h** Representative image of mock-infected (**g**) and OROV-infected (**h**) lungs, blue and green arrow indicated mixed interstitial and focal intra-alveolar infiltrate. **i-j** Representative image of mock-infected (**i**) and OROV-infected (**j**) heart, green arrow indicated mixed inflammatory cell population infiltration. **k-n** Representative images of mock-infected (**k**,**m**) and OROV-infected (**l**,**n**) liver, orange star and green arrow indicated hemorrhages and oil red O positive staining fat droplets. Scale bar = 100 µm. (mock, n = 3 and OROV, n = 2).

### Congenital OROV infection causes severe brain defects and extensive neural cell loss

Since OROV most severely affected fetal brains, we investigated the damages caused by this virus in more detail. To this end, we harvested embryonic brains at E17.5 (d5 pi) and performed cryo-sectioning followed by a nuclei counterstaining with DAPI, allowing visualization of the morphological alterations. Consistent with the intrauterine growth retardation, the cerebral hemispheres of OROV-infected brains were significantly reduced in size compared to the mock-infected ones at the same developmental stage (Fig. 3a-c). Although OROV-infected brains had smaller hemispheres, they showed a significant enlargement of the ventricles in comparison to mock-infected brains (Fig. 3d). Also, while the structure of the cerebral cortex in OROV-infected brains appeared similar to that in mock brains, featuring a progenitor zone, an intermediate zone and a cortical plate (Fig. 3a, b), significant differences were noted in the cerebral cortex. OROV-infected brains displayed a microcephaly-like phenotype, characterized by a lower number of DAPI-positive cells, a reduced cell density in the cortex as well as thinner cortices (Fig. 3e-g). Given the neurotropic nature of OROV, we performed additional immunostainings targeting markers of different neural brain cell types to determine which of these cells were affected the most in the OROV-infected brain. More specifically, to stain the progenitor neural cells of the cerebral cortex and striatum we used an antibody targeting the SOX2 marker^9^. To stain the projection neurons populating the deep-layer of the cortex as well as the projection medium spiny neurons in the striatum, we used an anti-CTIP2 antibody, whereas an antibody targeting the SATB2 marker was used to stain the neurons in the upper cortical layer^11^.

**Fig. 3:**
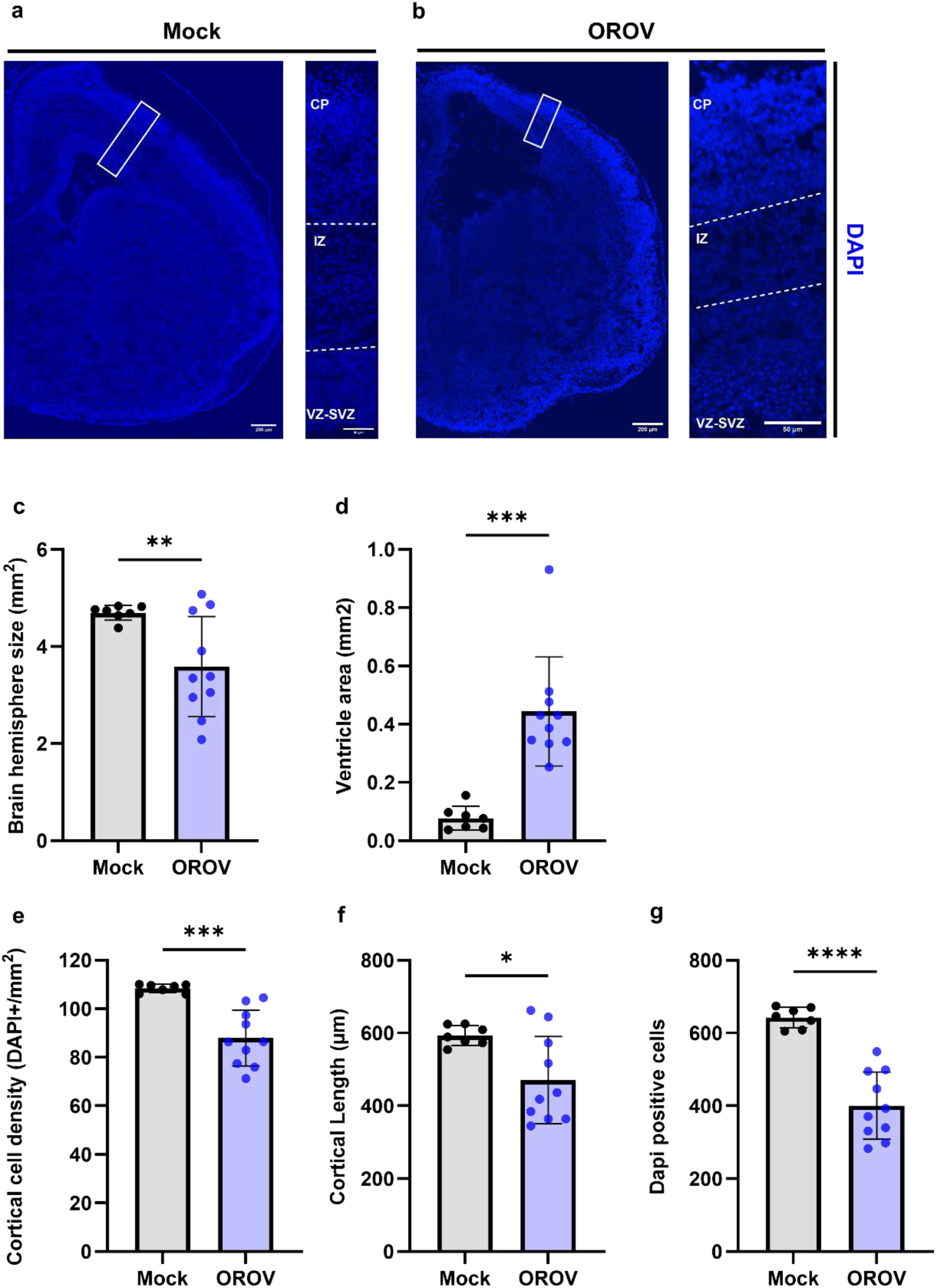
Congenital OROV infection causes brain abnormalities in embryos. **a**-**b** Representative images of E17.5 embryonic mouse brain after mock-infection (n = 7)(**a**) and OROV-infection (**b**) (n = 10). Reduction of the brain size was assessed by measuring the brain hemisphere size (**c**). Ventriculomegaly was estimated by measuring the ventricle area (**d**) Microcephaly-like phenotypes were assessed by measuring the cortical cell density (**e**), the cortical length (**f**) and the number of DAPI-positive cells (**g**). The scale bars in (**a**) and (**b**) represent 200 µm and 50 µm in the enlargements. In (**c**–**g**), data are presented as mean ± standard deviation. Statistical significance of differences was determined by the unpaired t test with Welch’s correction (**c**-**g**). Only statistically significant differences are shown (****p < 0.0001;***p<0.001; **p < 0.01; *p<0.05).

Compared to mock-infected brains, we observed a significant reduction in the SOX2 expression marker in the cortical progenitor region of OROV-infected brains (Fig. 4a, f, g). The percentage CTIP2-positive cells in OROV-infected brains, however, was comparable to that in mock-infected brains (Fig. 4b), albeit with a more compact distribution (Fig. 4h, i). Moreover, the upper cortical layer had (mostly) disappeared in the majority (∼75%) of the OROV-infected brains, characterized by a dramatic reduction of SATB2-positive cells (Fig. 4c, j, k). By contrast, the striatum of OROV-infected brains contained a lower percentage of both SOX2- and CTIP2-expressing cells (Fig. 4d, e, l, m) compared to mock-infected brains. Along with the decrease in CTIP2-positive cells and/or SOX2-expressing cells in the striatum and cortex, massive apoptosis in these regions was observed, as illustrated by a substantial increase in anti-cleaved caspase 3 (ACC3) expression (Fig. 4n, o). Although differences in severity were noted in OROV-infected brains, with some brains presenting a less dramatically altered phenotype, both moderately and severely affected brains displayed abundant apoptosis in the striatum, while the ACC3 expression was only minor in the cerebral cortex of moderately affected brains (Fig. 4p, q). Regardless of the severity of the brain damage, the ACC3 signal was always present in the progenitor zone of the striatum and the cerebral cortex (Fig. 4p, q). In conclusion, congenital OROV infection of the brain led to a decrease in the number of SOX2-positive cells in both the striatum and the cortex, while the number of CTIP2-positive cells was only reduced in the striatum. This was accompanied by massive apoptosis, particularly in the striatum.

**Fig. 4:**
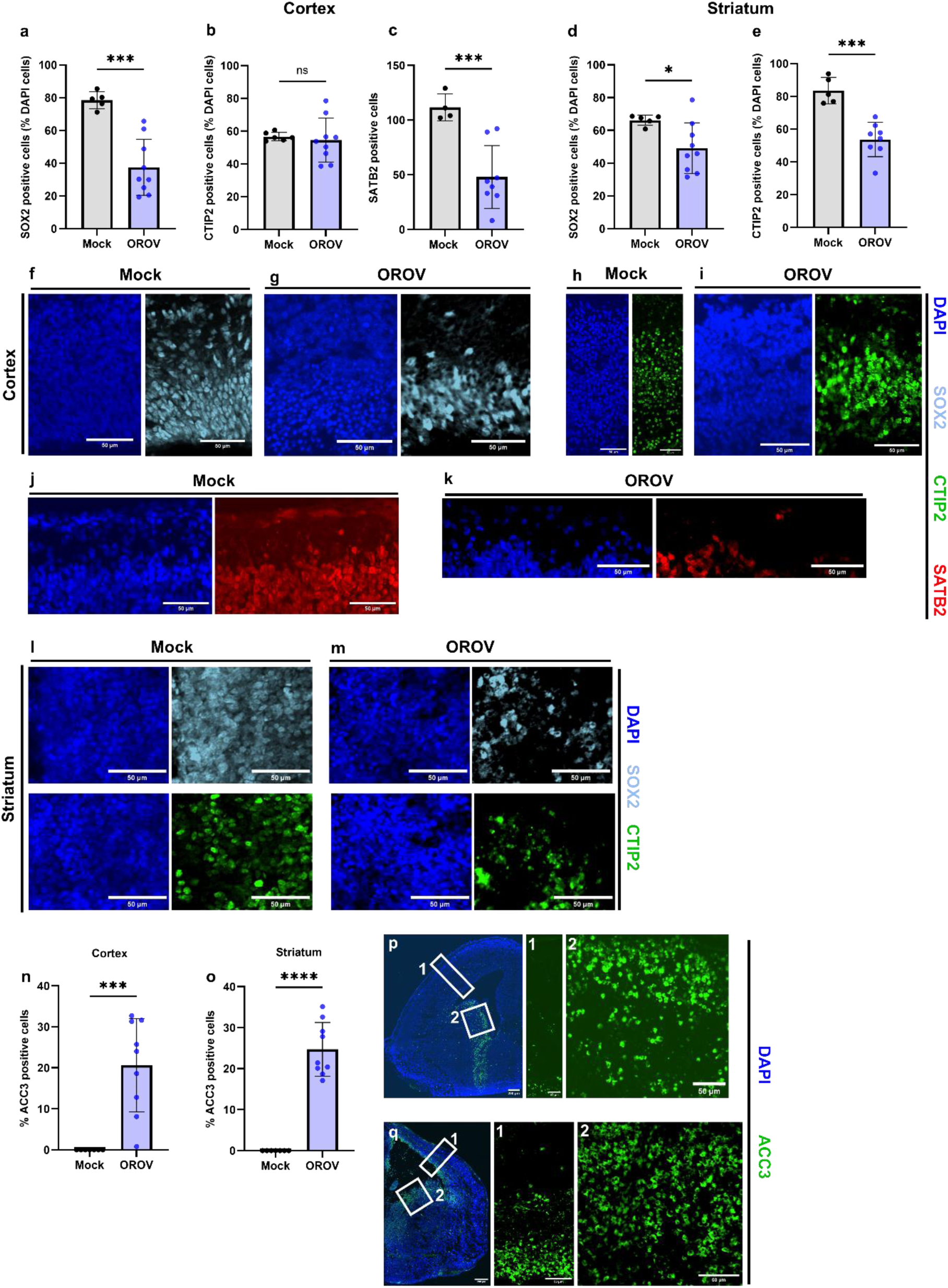
Impact of OROV congenital infection on neural cell populations and brain architecture. **a-c** Measurement of the number of SOX2-positive cells (**a**), CTIP2-positive cells (**b**) and SATB2-positive cells (**c**) in the cerebral cortex in mock-infected (n = 5) and OROV-infected brains (n = 9). **d,e** Measurement of number of SOX2-positive cells (**d**) and CTIP2-positive cells (**e**) in the striatum in mock-infected (n = 5) and OROV-infected brains (n = 8). **f**-**k** Representative images of E17.5 cerebral cortex of mock-infected (**f,h,j**) and OROV-infected brains with a severe phenotype (**g,i,k**). Dark blue, light blue, green and red indicated DAPI, SOX2, CTIP2 and SATB2 staining, respectively. **l,m** Representative images of E17.5 striatum of mock-infected (**l**) and OROV-infected with a severe phenotype (**m**). Dark blue, light blue, green and red indicated DAPI, SOX2, CTIP2 and SATB2 staining, respectively. **n**,**o** Measurement of the number of anti-cleaved caspase3 (ACC3) positive cells at E17.5 in the cerebral cortex (**n**) and striatum (**o**) of OROV-infected brains. **o**,**p** Representative images of a moderately affected (**p**) and severely affected (q) OROV-infected brains. Dark blue and green indicated DAPI and ACC3 staining, respectively. Statistical significance of differences was determined by the unpaired t test with Welch’s correction. Statistically significant differences are shown (****p < 0.0001;***p<0.001; *p<0.05), ns = not significant. The scale bars represent 50 µm.

### A pronounced pro-inflammatory immune response in OROV-infected embryonic brains

To investigate whether and to what extent OROV had triggered an immunological response, we measured the levels of a small selection of cytokines and chemokines (IL-6, IL-1β, TNF-α, IFN-γ and CCL5) in various embryonic organs (brain, lungs, heart, and liver) collected at E17.5. All organs studied had elevated expression levels of the pro-inflammatory cytokines IL-6, IL-1β, TNF-α and IFN-γ, and the pro-inflammatory chemokine CCL5 compared to mock-infected embryonic organs (Fig. 5a). Interestingly, of all studied organs in the embryonic mice, the pro-inflammatory immune response was the strongest in the brain (Fig. 5a), which corresponded with the higher viral RNA levels in the brain than in the other organs (Fig. 1d). This finding together with our earlier observation that OROV-infected embryonic brains were heavily infiltrated with neutrophils (Supplementary Fig. 2b), prompted us to more thoroughly investigate the neutrophil infiltrations in these brains. We therefore conducted additional immunostainings with the Ly-6G monoclonal antibody, which specifically targets the neutrophil populations. In addition, we also further characterized the microglial cells (Iba1 staining), which were earlier found to also be present in OROV-infected brains (Supplementary Fig. 2b). Whereas neutrophils were not observed in mock-infected brains (Supplementary Fig. 4), both the striatum and the cerebral cortex of OROV-infected brains showed a massive infiltration of neutrophils (Fig. 5b, c), although the number of neutrophils appeared to be higher in the striatum than in the cortex (Fig. 5d, e). Further characterization of the microglial cells revealed that the microglia in OROV-infected brains presented an activated phenotype with a round shape compared to those in mock-infected brains (Fig. 5 f-h). Also, OROV-infected brains presented some microglial nodules, which are aggregations of activated microglial cells (Fig. 5h, right panel). Taken together, we show that congenital OROV infection induced a pro-inflammatory immune response in all embryonic organs studied, as demonstrated by increased expression levels of pro-inflammatory cytokines and chemokines. Moreover, the inflammatory response was the strongest in embryonic brains (which also had higher viral RNA levels), as evidenced by a greater number of activated microglial cells and a pronounced recruitment of neutrophils.

**Fig. 5:**
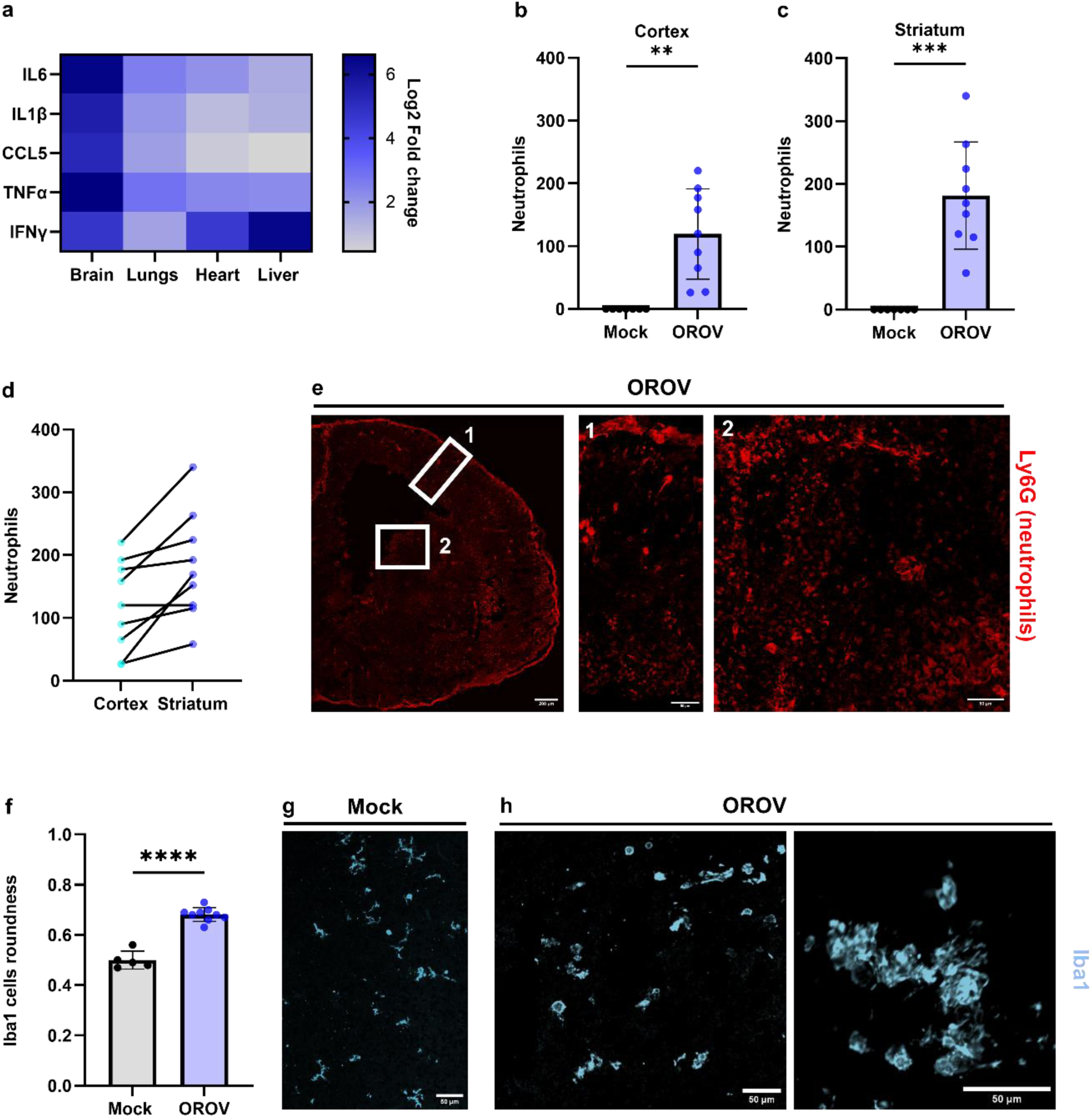
Congenital OROV infection leads to an inflammatory response within the embryonic organs. **a** Heatmap of expression profiles (log2 fold change) of the cytokines IL6, IL1β, TNFα, IFNγ and the chemokine CCL5 in the embryonic brain, lungs, heart and liver of OROV-infected embryos compared to mock-infected embryos at E17.5. **b**,**c** Measurements of the number of neutrophils in the cerebral cortex (**b**) and striatum (**c**) of OROV-infected brains at E17.5. **d** Comparison of the number of neutrophils between the cortex and striatum of OROV-infected brains. **e** Representative images of E17.5 OROV-infected brains stained by immunofluorescence for neutrophils (Ly6G). **f** Measurement of Iba1 cells roundness in mock-infected and OROV-infected brains. **g**,**h** Representative images of microglial cells (iba1) in mock-infected (**g**) and OROV-infected (**h**) brains. Statistical significance of differences was determined by the unpaired t test with Welch’s correction (**f**) Statistically significant differences are shown (***p<0.001; **p<0.01). The scale bars represent 200 µm and 50 µm.

## Discussion

OROV circulated for decades in Central and South America, and was historically associated with self-limiting febrile illness or asymptomatic infections and causing only sporadic epidemic events. Consequently, OROV was previously not regarded as a significant threat to human health. The (still ongoing) 2023-2024 outbreak in Latin America has reshaped this perception. For the first time, severe and fatal cases have been documented in young and previously healthy individuals without comorbidities. Worryingly, recent PAHO reports describe cases of OROV vertical transmission, some resulting in miscarriage. Moreover, suspected cases of OROV vertical transmission were linked to congenital disease with severe brain abnormalities reminiscent of congenital Zika syndrome (CZS), including microcephaly, ventriculomegaly and brain calcifications. Whereas vertical transmission of OROV has been confirmed clinically, proof of a causal link between *in utero* OROV infection and congenital disease lagged. Given the expanding geographical spread of OROV, with the emergence in previously unaffected countries including imported cases in North America and Europe, there is an urgent need to study the consequences of vertical OROV transmission for fetal development.

Animal models have been instrumental in the investigation of viral pathogenesis. OROV pathogenesis has been studied in three-week-old Syrian hamsters^12^ and suckling mice^13^. Wildtype (C57BL/6, CD1) adult mice were not suitable as OROV was unable to replicate efficiently and because OROV infections did not cause overt pathology in these mice due to the robust RIG-I-like receptor signaling pathway, downstream regulatory transcription factors (IRF-3 / IRF-7), and the type 1 interferon (IFN) response^14^. Consequently, studies relied on IFN deficient mice to permit viral replication. Recently, however, efficient OROV replication and vertical transmission was demonstrated in WT C57BL/6 mice^15^. Yet, this model cannot be used to study OROV-induced congenital disease because of the absence of overt pathology in the mice, and suckling mice are unsuitable for studying congenital brain alterations caused by OROV, as their cerebral cortex is already largely formed and neurodevelopmental mechanisms during embryonic development are different from those present at birth. To address this gap, we adapted the intraplacental (IPL) infection model previously developed for Zika virus^7^, delivering OROV directly into the labyrinth (fetal part) of the placenta in WT mice.

In human cases of congenital OROV infection, the virus has been detected in multiple fetal organs (i.e. heart, lungs, liver, spleen and brain) with histopathology revealing widespread lesions in the infected organs and severe brain involvement in particular^16^. Consistently, IPL infection with OROV in our model, led to intrauterine growth retardation, characterized by a reduced crown-rump length and a lower body weight of the embryonic mice. Robust viral replication was detected in all major organs studied (heart, lungs, liver and brain), with the highest viral RNA levels in the brain. Histopathological examination of the infected organs revealed widespread abnormalities, including hepatic steatosis and pneumonitis, with the brain and the spinal cord showing the most severe defects. Neuropathological hallmarks included hemorrhages, neural loss, and immune cells infiltration, closely mirroring findings from autopsied OROV-infected babies^6,16^. Some mouse embryos also exhibited subcutaneous edema and/or hydrops, again paralleling reported human cases. Given the striking neurotropism of OROV, we investigated whether our model recapitulates the spectrum of congenital brain malformations reported in humans. Indeed, OROV-infected embryonic brains had smaller hemispheres than the mock-infected counterparts at the same developmental stage. In addition, OROV-infected embryonic brains presented with ventriculomegaly and a microcephaly-like phenotype (i.e. cortical thinning with reduced neural density), indicating that OROV infection directly disrupts fetal brain development and that the brain abnormalities largely correspond to those observed in human cases. Furthermore, we demonstrated that the development of brain defects directly correlates to an *in utero* infection by OROV.

Because the hallmarks of congenital OROV infection included microcephaly and ventriculomegaly, we next focused on the specific impact of OROV infection on the neural populations of two major regions of the telencephalon: the cerebral cortex and the striatum, both of which are essential for higher-order brain functions (e.g. language, learning, cognition, amongst others). The cortex is a highly organized structure consisting of six neuron layers^17^, whereas the striatum is a less highly defined structure compared to the cortex^18^. Perturbations in the development of either region are associated with lifelong neurological and psychiatric disorders. Our analyses revealed profound alterations in the neural populations of the cortex and striatum. In both the cortex and striatum of OROV-infected brains, the SOX2 population, representing the progenitor cells^9^, were markedly reduced. In addition, although the percentage of the CTIP2-positive cell population (representing the projection neurons populating the deep layer in the cortex^10^) appeared preserved, the distribution of these neurons was denser, reflecting impaired laminar organization. Lastly, SATB2-positive neurons (representing the projection neurons in the upper cortical layer^11^) were drastically reduced or nearly absent. In the striatum, both SOX2-positive progenitors and CTIP2-positive spiny projection neurons^19^ were markedly decreased, pointing to a collapse of striatal neurogenesis and circuitry. Consistent with these findings, apoptotic activity was strongly elevated in the progenitor zones of both the cerebral cortex and the striatum, hence correlating with the severe reduction in the SOX2-positive populations.

Together, these results show that congenital OROV infection in our model can cause devastating neural loss and structural disorganization in both the cerebral cortex and the striatum. Interestingly, loss of SOX2 progenitor cells has been linked to enlargement of the ventricles in mice^20^, thus corroborating the ventriculomegaly induced by OROV in our model. Even more, the compact morphology of the CTIP2 cortical layer and the almost complete absence of the upper layer are in line with the microcephalic phenotype observed in OROV-infected human fetuses. We further assessed the immunopathological response to congenital OROV infection in our model. All infected organs displayed pro-inflammatory cytokine upregulation, with the brain showing the strongest inflammatory response, which thus appears to be linked to the higher viral load measured in the brain. OROV infection triggered a massive influx of neutrophils into the brain, which can be attributed to the higher levels of IL-6, IL-1β and TNF-α; all strong neutrophil attractants^21^. Moreover, neutrophil recruitment was noted in both the striatum and the cerebral cortex, although numbers tended to be higher in the striatum. This correlated with the sites of apoptosis in both regions, but with a higher prevalence in the striatum. Microglia activation and formation of microglial nodules were also evident, reflecting findings in autopsied human cases. Whereas neutrophils can have a protective role in some viral infections by contributing to viral clearance^22,23^, excessive recruitment has been implicated in exacerbating neuropathology in other viral infections^24^, raising the possibility that OROV-induced neuroinflammation contributes to fetal brain damage. The precise role of neutrophils in congenital OROV infection -protective, detrimental, or both-warrants further investigation.

In conclusion, our findings establish the IPL mouse model as a highly relevant model to study the consequences of congenital OROV infection for the developing fetus as it recapitulates key clinical features observed in human congenital OROV cases, including multi-organ infection, severe neuropathology, and hallmark developmental defects such as microcephaly and ventriculomegaly. These findings suggest that congenital OROV infection can lead to miscarriage, neonatal death, or long-term neurodevelopmental disorders, alike congenital ZIKV infection. Recent complementary studies reinforce this conclusion: OROV was shown to be vertically transmitted in immunocompetent mice^14^, infect human trophoblasts and placenta explants^25^, and disrupt brain development in brain organoids^26^. Altogether, these results firmly position OROV as an emerging public health threat, in particular for pregnant women and their developing fetus. Enhanced surveillance of OROV circulation, systematic screening of pregnant women during outbreaks, and close clinical monitoring of pregnancies in cases of a suspected OROV infection are therefore vital to mitigate the potential impact of a congenital OROV infection.

## Methods

### Cells

VeroE6 cells (African green monkey kidney cells; Vero 76, clone E6; ATCC: Cat# CRL-1586) were cultured in DMEM supplemented with 10% fetal bovine serum (FBS; HyClone) and 2 mM glutamine (Thermo FisherScientific). BHK-21J (baby hamster kidney fibroblasts; provided by Dr. Peter Bredenbeek at the Leiden University Medical Center (LUMC), the Netherlands) were cultured in DMEM supplemented with 10% FBS (HyClone) VeroE6 and BHK21J cells were incubated at 37 °C in presence of 5% CO_2_. VeroE6 and BHK21J cells were regularly tested for mycoplasma contamination.

### Oropouche virus

The Oropouche virus (OROV) strain OROV_IRCCS-SCDC_1/2024_ was isolated from a semen sample of a patient returning from Cuba and admitted to the IRCCS Sacro Cuore Don Calabria Hospital in Negrar di Valpolicella (Italy) in June 2024 and kindly provided by Dr. Concetta Castilletti ^8^. The whole genome sequence is available in GenBank (BankIt2843215; S segment: PP952117, M segment: PP952118, L segment: PP952119). The isolate was propagated on VeroE6 cells to generate the virus stock. The infectious virus titer (expressed as plaque forming units [PFU] per ml) was determined by plaque assay, as described previously^27^.

### Mouse experiments

Pregnant SWISS mice were purchased from Janvier Labs. Mice were housed at KU Leuven at biosafety level 3 in individually ventilated cages (type GM500, Sealsafe Plus, Tecniplast) at 21°C, 55% humidity and 12:12 light/dark cycles. Animals were provided with food and water ad libitum as well as with cardboard play tunnels and cotton/tissues as extra bedding material. Allocation to experimental groups was performed randomly. Housing conditions and experimental procedures were approved by the ethical committee of KU Leuven (license DMIT-190/2024 and DMIT-032/2025) following institutional guidelines approved by the Federation of European Laboratory Animal Science Associations (FELASA).

### Intraplacental infection

Time-mated SWISS mice, 8–12 weeks of age, were used for the OROV vertical transmission experiments. All surgeries on wild-type female SWISS mice were performed at the same time, with embryonic day (E) 0.5 corresponding to the afternoon following the day of mating. At E12.5, preoperative analgesia (0.1 mg/kg of buprenorphine; Ethiqa XR) was administered subcutaneously before induction of anesthesia. The pregnant females were anesthetized using isoflurane (Abbot Laboratories Ltd.), after which the mice were shaved. Next, a small incision (1.0-1.5 cm) was made through the lower ventral peritoneum, followed by the careful extraction of the uterine horns, which were then carefully placed on warm humidified gauze pads. The developing embryos were challenged intra-placentally with 10-20 PFU OROV or PBS (‘mock’). After infection, incisions were sutured and disinfected. Mice were maintained on a heat pad during the whole procedure. Four to five hours post-surgery, mice were checked for any signs of abortion.

### RNA isolation and RT-qPCR

Total RNA was isolated from embryonic tissue using Trizol (Ambion, Life Technologies), according to the manufacturer’s protocol. RNA was eluted in 30 µl of RNAse free water. Quantification of OROV genome and cytokines/chemokines copy numbers was performed on a QuantStudio5 platform (Applied Biosystems) by quantitative reverse transcription PCR (RT-qPCR) using the one step RT-qPCR iTaq Universal probe kit (Ref. 1725141, Bio-Rad). Primer and probe sequences for OROV RNA and cytokine/chemokine quantification are provided in Supplementary Table 1. For OROV, Ct values were converted into a relative number of OROV RNA copies/μl using the formula y = a*ln(x) + b, where a is the slope of the standard curve, b is the y-intercept of the standard curve and y is the Ct value for a specific sample. For cytokines measurement, results are expressed as Log2 fold change = 2^-ΔΔCt^, where 2^-ΔΔCt^ = [(Ct gene of interest – Ct housekeeping gene)infected_sample – (Ct gene of interest – Ct housekeeping gene)mock_sample)]. The sequence of the cytokine primers and probes were adopted from Overbergh and colleagues^28,29^.

### Cryo-sectioning

Embryonic mouse heads and bodies were harvested and fixed overnight at 4 °C in 4% paraformaldehyde (PFA). Brains were dissected in 0.1 M PBS (pH 7.4). Dissected embryonic brains and bodies were cryoprotected by immersion in a sucrose gradient (first 15% sucrose in PBS followed by 30% sucrose in PBS; both overnight at 4 °C). Samples were then embedded in Polyfreeze (Sigma-Aldrich) or Epredia™ Neg-50™ Frozen Section Medium (Fisher Scientific, 12678086). Sections of 14 μm were obtained by cryo-sectioning (Leica) onto slides (Epredia SuperFrost Plus™ Adhesion slides, Fisher Scientific).

### Immunofluorescence

For immunofluorescence of brain and body sections, antigen retrieval was performed at 95 °C for 15 min before incubation with primary antibodies. Slides were then incubated in blocking solution (Normal Donkey Serum 10%, Bioconnect Life Sciences) for 1 h at room temperature. Sections were incubated with primary antibodies (Supplementary Table 2) overnight at 4 °C. The following day, slides were stained using the secondary antibodies (Supplementary Table 2) and nuclei were counterstained with DAPI (1:1000) for 2 h at room temperature. Slides were mounted in Dako Fluorescence Mounting Medium (Agilent).

### Hematoxylin and Eosin staining

Brains and whole embryonic body sections were stained for Hematoxylin and Eosin using the H&E Staining Kit (Hematoxylin and Eosin, Abcam, ab245880) following the manufacturer’s protocol.

### Confocal microscopy and analysis

Confocal images of sections of embryonic mouse brains and whole bodies were acquired using a Leica DMi8 confocal spinning disc microscope at a 25X magnification. Image analysis and processing were performed with ImageJ 1.42q 276 (Wayne Rasband, National Institutes of Health), Fiji (v2.0.0-rc-54/ 1.51 h; https://imagej.net/Fiji) or Cellpose^30^ software packages.

### Statistics

Statistical analyses were performed using GraphPad Prism v9.3.1. Results were first tested for normality using the Shapiro-Wilk test. In case of a normal distribution of the data, an unpaired t-test with Welch’s correction was performed. When the data did not follow a normal distribution, a Mann-Whitney test was performed. Differences were considered statistically significant when p < 0.05.

## Acknowledgments

We thank the colleagues at the Rega animal facility at KU Leuven for technical assistance. This work was funded by the DURABLE project (to J.N.). The DURABLE project has been co-funded by the European Union, under the EU4Health Programme (EU4H), project no. 101102733. The funders had no role in the study design, data collection and interpretation, or the decision to submit the work for publication.

## Author Contributions

M.D.: Conceptualization, planning, coordination and execution of the work. R.M.V.: Histopathology pictures acquisition and interpretation. N.C., M.F., S.H., Technical assistance. A.H.: Image analysis and analysis software assistance. S.J.F.K. and J.N.: Securing of funding, conceptualization. S.J.F.K. and M.D. wrote the manuscript with contributions from J.N., and comments from all authors.

## Competing Interests

The authors declare no competing interests.

## Data and Materials Availability

All data generated and analyzed in this paper are included in this article. Additional imaging data related to the immunostainings or the hematoxylin and eosin staining are available from the corresponding authors upon request. Source data are provided with this paper.

---

**Supplementary Table 1:**
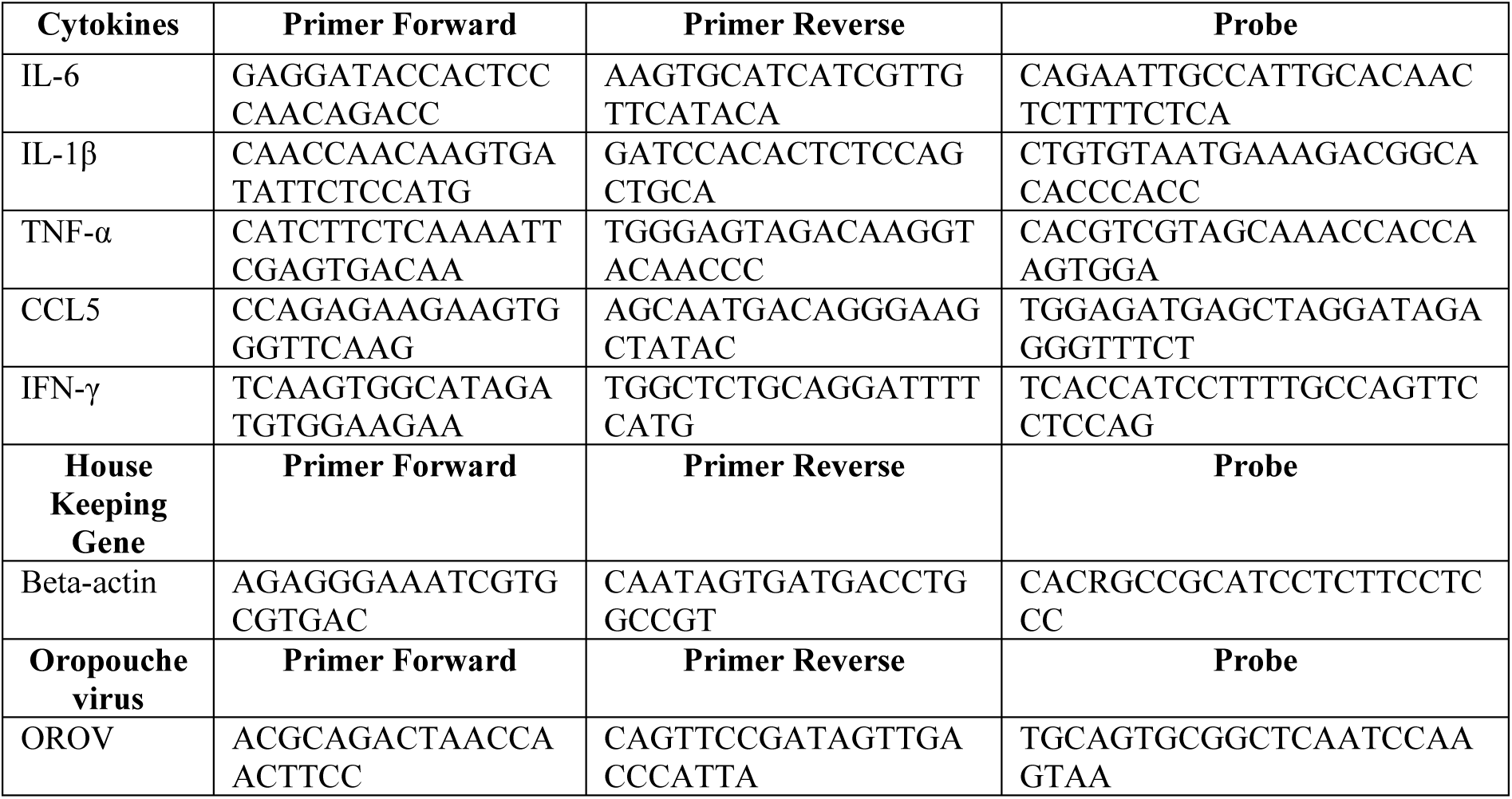
Primers and Probes included in this study.

**Supplementary Table 2:** List of antibodies used in this study.

| Primary antibodies | Catalog Number | Company | Concentration | Antigen retrieval |
| --- | --- | --- | --- | --- |
| Cleaved Caspase-3 (Asp175) Antibody | 9661 | Cell Signaling Technology | 1:300 | No |
| Anti-SATB2 antibody [EPNCIR130A] | ab92446 | Abcam | 1:500 | Yes |
| Ly-6G Monoclonal Antibody (1A8-Ly6g) | 16-9668-82 | ThermoFisher Scientific | 1:100 | No |
| CUX1 Polyclonal antibody | 11733-1-AP | Proteintech | 1:200 | Yes |
| Anti-Iba1 antibody | ab5076 | Abcam | 1:300 | No |
| Anti-Ctip2 antibody [25B6] | ab18465 | Abcam | 1:500 | Yes |
| Human/Mouse/Rat SOX2 Antibody | AF2018 | Bio-Techne | 1:500 | Yes |
| Recombinant Anti-CCR2 antibody | ab273050 | Abcam | 1:200 | Yes |
| Secondary antibodies | Catalog Number | Company | Concentration |  |
| Alexa Fluor® 488-AffiniPure Donkey Anti-Rat IgG (H+L) | 712-545-153 | Jackson ImmunoResearch | 1:1000 | NA |
| Donkey anti-Mouse IgG (H+L) Alexa Fluor 555 | A-31570 | Thermo Scientific | 1:1000 | NA |
| Donkey anti-Goat IgG (H+L) Alexa Fluor 647 | A-21447 | Thermo Scientific | 1:1000 | NA |
| Donkey anti Rabbit IgG (H+L) Alexa Fluor 488 | A-21206 | Thermo Scientific | 1:1000 | NA |

## Supporting information

**Supplementary Fig. 1.**
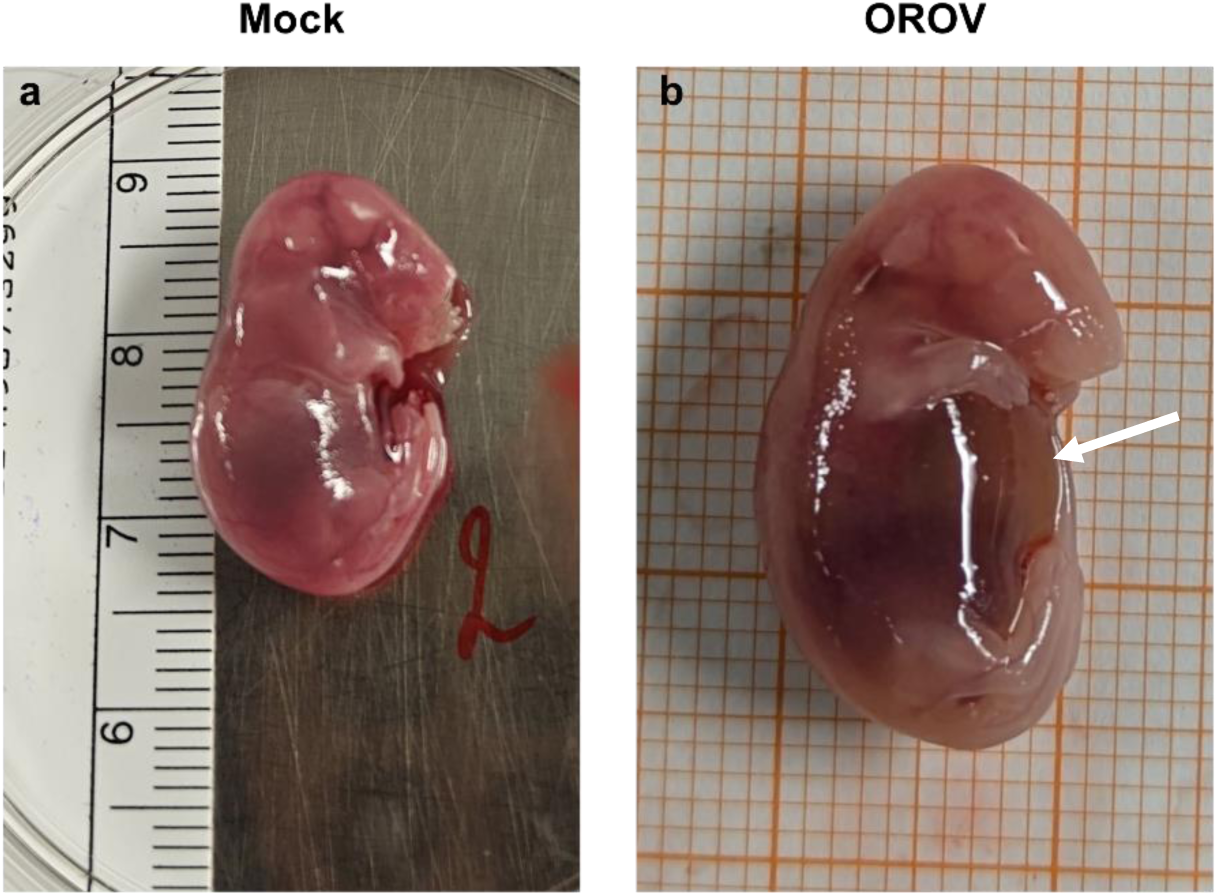
Congenital OROV infection can cause subcutaneous edema and hydrops in mouse embryos. **a**. Representative picture of a mock-infected embryo at E17.5. **b**. Representative picture of an OROV-infected embryo at E17.5 (d5pi).

**Supplementary Fig. 2.**
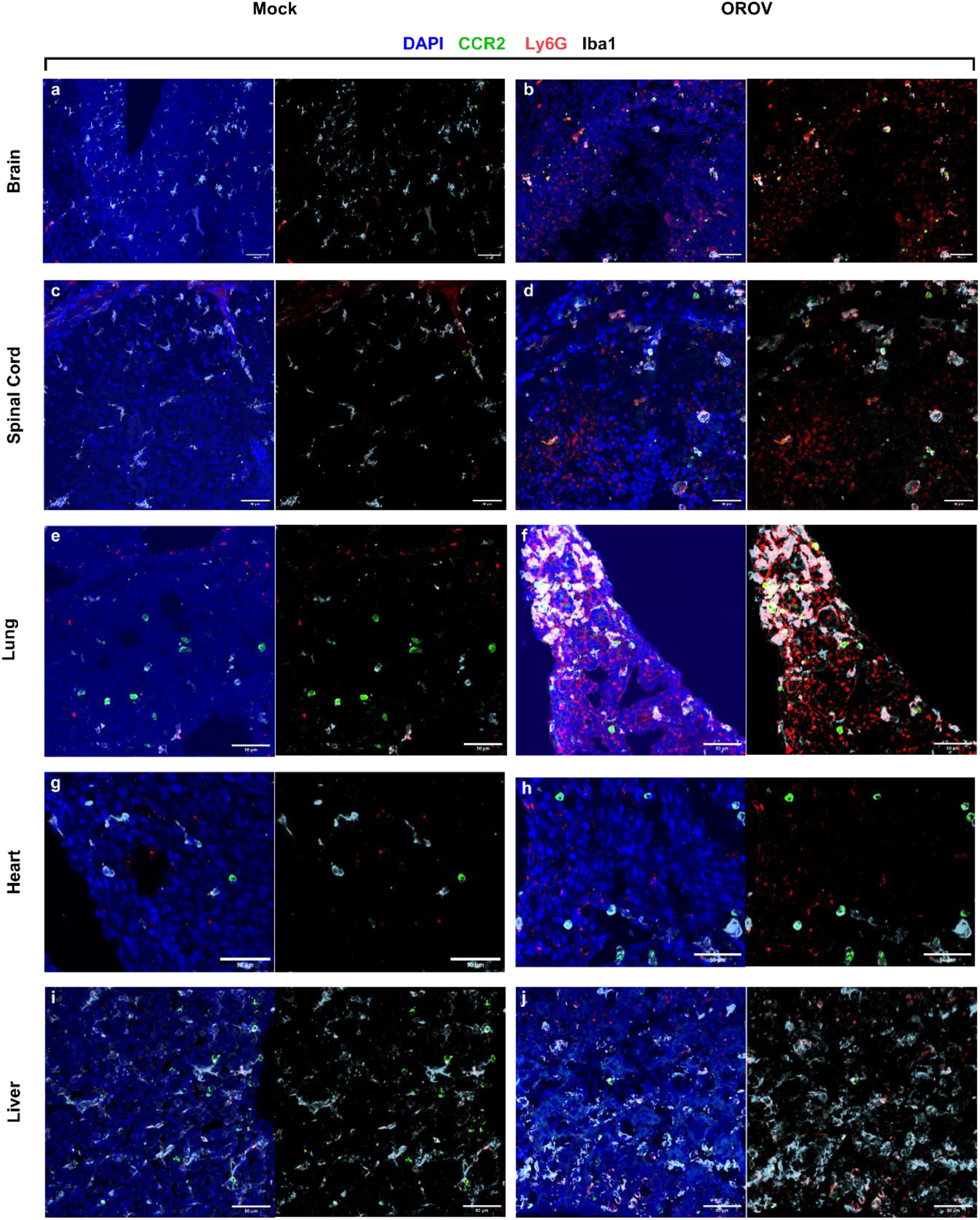
Distinct immune marker expression in embryonic tissues following OROV infection. **a,j.** Representative images of E17.5 mock-infected brain (**a**), spinal cord (**c**), lung (**e**), heart (**g**) and liver (**i**), and OROV-infected brain (**b**), spinal cord (**d**), lung (**f**), heart (**h**) and liver (**j**). Dark blue, green, red and white represent DAPI, CCR2, Ly6G and Iba1 staining, respectively. The scale bars represent 200 µm.

**Supplementary Fig. 3.**
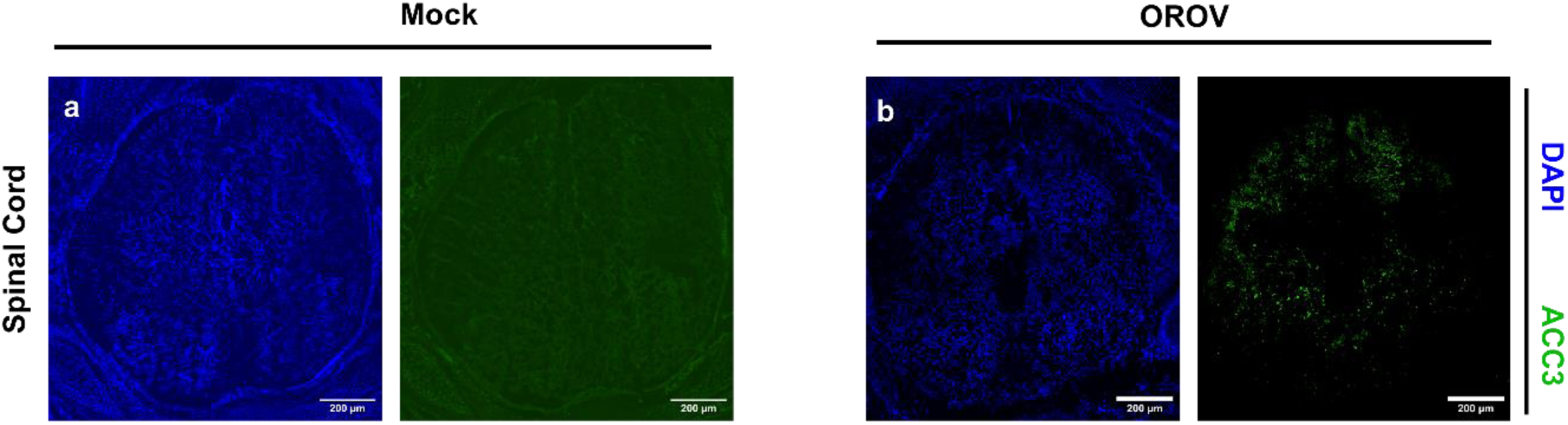
OROV infection induces ACC3 expression in the embryonic spinal cord. **a,b.** Representative images of E17.5 mock-infected spinal cord (**a**)and OROV-infected spinal cord (**b**). Dark blue, green indicated DAPI and ACC3 staining, respectively. The scale bars represent 200 µm.

**Supplementary Fig. 4.**
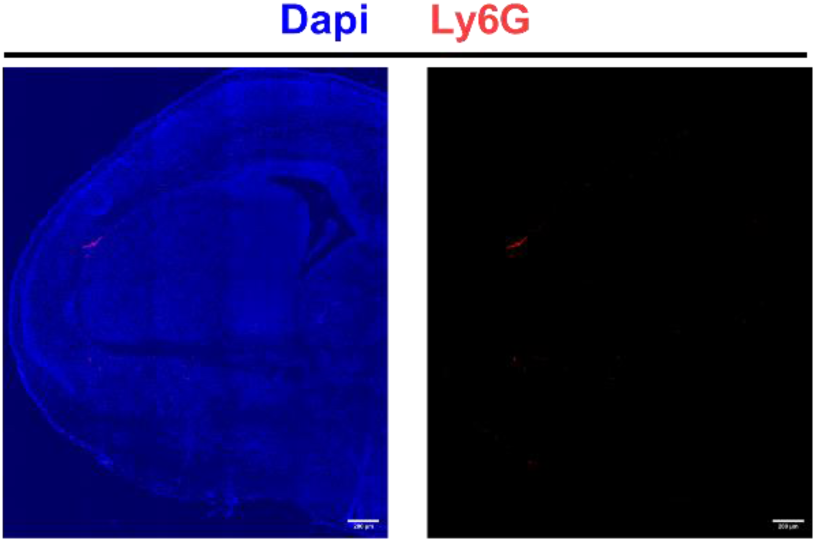
Absence of neutrophils in a E17.5 mock-infected brain. Representative picture of a E17.5 mock-infected brain. Dark blue and red indicated DAPI and Ly6G staining respectively. The scale bars represent 200µm

## Notes

### Competing Interest Statement

The authors have declared no competing interest.

